# Impact of experimental bias on compositional analysis of microbiome data

**DOI:** 10.1101/2023.02.08.527766

**Authors:** Yingtian Hu, Glen A. Satten, Yi-Juan Hu

## Abstract

Microbiome data are subject to experimental bias that is caused by DNA extraction, PCR amplification among other sources, but this important feature is often ignored when developing statistical methods for analyzing microbiome data. McLaren, Willis and Callahan (2019) proposed a model for how such bias affects the observed taxonomic profiles, which assumes main effects of bias without taxon-taxon interactions. Our newly developed method, LOCOM (logistic regression for compositional analysis) for testing differential abundance of taxa, is the first method that accounted for experimental bias and is robust to the main effect biases. However, there is also evidence for taxon-taxon interactions. In this report, we formulated a model for interaction biases and used simulations based on this model to evaluate the impact of interaction biases on the performance of LOCOM as well as other available compositional analysis methods. Our simulation results indicated that LOCOM remained robust to a reasonable range of interaction biases. The other methods tended to have inflated FDR even when there were only main effect biases. LOCOM maintained the highest sensitivity even when the other methods cannot control the FDR. We thus conclude that LOCOM outperforms the other methods for compositional analysis of microbiome data considered here.

## Introduction

Experimental bias introduced during DNA extraction and PCR amplification, among other steps in the experimental pipeline, is a pervasive feature of microbiome data. Absent experimental procedures that produce unbiased data, it is therefore necessary to account for experimental bias when analyzing microbiome data. Fortunately, McLaren, Willis and Callahan (MWC) [1] have proposed a model to explain how experimental bias affects microbiome data in which the observed relative abundance of each taxon is a product of the taxon’s true relative abundance and a taxon-specific bias factor, normalized over all taxa observed in the sample. Each taxon-specific bias factor represents the accumulation of multiplicative biases over all the steps in the experimental pipeline, so that multiple sources of bias are described by a single factor. With the MWC model in mind, we developed LOCOM [2], a logistic-regression-based compositional analysis method for detecting differentially abundant taxa, that only estimates parameters that are free from bias, i.e. are not affected by bias factors. To the best of our knowledge, LOCOM was the first method that accounted for experimental bias and was shown both analytically and numerically to be fully robust to any bias that follows the MWC model; none of the other existing methods for testing taxon differential abundance have considered experimental bias.

However, the MWC model assumes no between-taxon “interaction” bias, i.e., the presence or abundance of one taxon does not affect the bias factors of any other taxa in the sample. LOCOM may not be bias-robust if the “main effect” biases of the MWC model were supplemented by interaction biases. In evaluating the MWC model, Zhao and Satten [3] in fact found a small amount of evidence for interaction biases, although with smaller magnitude than the main effect biases in the MWC model; this finding is the motivation for this study. Here, we formulate a model for interaction biases and use simulations based on this model to evaluate the impact of interaction biases on the performance of LOCOM, as well as a number of existing compositional analysis approaches (ANCOM [4], ANCOM-BC [5], fastANCOM [6], ALDEx2 [7], WRENCH [8], DACOMP (v1.26) [9], LinDA [10], and the Wilcoxon rank-sum test of log-ratio transformed data after adding pseudocounts of either 0.5 or 1). Thus, this report provides the first assessment of the impact of interaction biases on analysis of microbiome data.

## Methods

We generalize the MWC model using the framework of Zhao and Satten [3] by adding an interaction bias *θ_jj′_* to the log-linear model that relates the expected value of the observed relative abundances and true relative abundances:

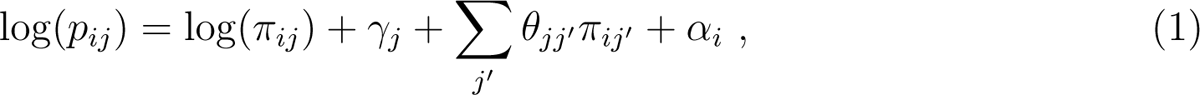

where *π_ij_* is the true relative abundance of taxon *j* in sample *i*, *p_ij_* is the expected value of the observed relative abundance, *γ_j_* is the main effect bias for taxon *j* in the MWC model, and *α_i_* is the sample-specific normalization factor that ensures the compositional constraint on *p_ij_*. We followed [3] to introduce the effects of covariates *X_i_* on taxon *j* by replacing log(*π_ij_*) in (1) by 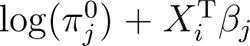, where *β_j_* contains the effect sizes and 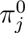 is the true relative abundance of taxon *j* when *X_i_* = 0. The interaction *θ_jj′_* determines the extent to which taxon *j*′ affects the bias factor for taxon *j*. The assumption underlying the MWC model is *θ_jj′_* = 0 for all *j* and *j*′. To generate interactive biases in our simulation studies, we additionally express *θ_jj′_* relative to *γ_j_* as

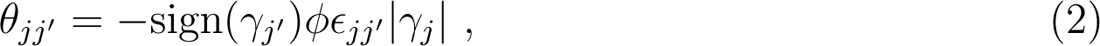

where *ϕ* is a constant for all taxa pairs that we call the magnitude of interaction biases, and *ϵ_jj′_* is a non-negative error term having mean one. The value 0.612 for *ϕ* and the sign in (2) reflect the findings of Zhao and Satten [3], who considered a model like (1) but with *π_ij′_* replaced by *I*[*π_ij′_* > 0] in the interaction term; more details are found in Supporting Information.

Our simulations for evaluating the effects of interaction biases on compositional analysis methods were based on models (1) and (2) and data on 856 taxa of the upper-respiratory-tract microbiome by Charlson et al. [11]. We considered both binary and continuous traits of interest without any confounding covariates; we also considered a binary confounder when the trait was binary. We used the two sets of causal taxa (i.e., taxa that are associated with the trait) that were used in [2], namely, a random sample of 20 taxa (referred to as M1) and the five most abundant taxa (M2); we assumed a common effect size *β* of the trait on all causal taxa. We designed three scenarios for the distribution of interaction biases: sampling the error term *ϵ_jj′_* non-differentially based on *N*(0.5, 0.1^2^) for all taxa (referred to as S-nondiff), differentially based on *N*(0.5, 0.1^2^) and *Beta*(0.5, 0.5) (a bimodal distribution with larger variance than 0.1^2^) for null and causal taxa, respectively (S-diff-causal), and differentially for a random selection of half of the taxa and the remaining taxa (S-diff-half). Unlike S-nondiff, S-diff-causal has modest (but trait-related) variation in *ϵ_jj′_* while S-diff-half has large variation in *ϵ_jj′_* that is not trait-related. Additional details of the simulation settings are found in Supporting Information, but generally follow the simulations in [2].

## Results

The empirical FDR and sensitivity of all aforementioned compositional analysis methods for detecting the causal taxa, for simulations with a binary trait and no confounder under scenarios M1 and M2, are displayed in Figures 1 and 2, respectively (results for simulations with confounders and continuous traits showed similar patterns and are therefore omitted here). Note that *ϕ* = 0 corresponds to no interaction bias at any taxa (i.e., main effect biases only) and *β* = 0 corresponds to no differential abundance at any taxa (i.e., the global null). In all cases, the FDR of LOCOM remained at or close to the nominal level as long as the magnitude of interaction bias *ϕ* < 1, which is substantially larger than the empirical value 0.612 observed in the Brooks data [3]; under the global null in particular, LOCOM always controlled the FDR regardless of the value of *ϕ*. It was only when both *ϕ* and *β* became unrealistically large that we observed moderate inflation in the FDR of LOCOM. The FDR inflation of LOCOM was similar in S-nondiff and S-diff-causal, because the interaction biases were similarly distributed at the majority of taxa which were null taxa; the inflation was (slightly) the largest in S-diff-half, because the interaction biases had the largest variability among the three scenarios at a given *ϕ*. The interaction biases caused some loss of sensitivity for LOCOM, but the drop was relatively small and LOCOM maintained the highest sensitivity among all methods in all of our simulation scenarios. The results of the other methods showed a similar trend to LOCOM with FDR increasing and sensitivity decreasing as *ϕ* increased, and increasing FDR inflation as *β* increased.

**Figure 1:**
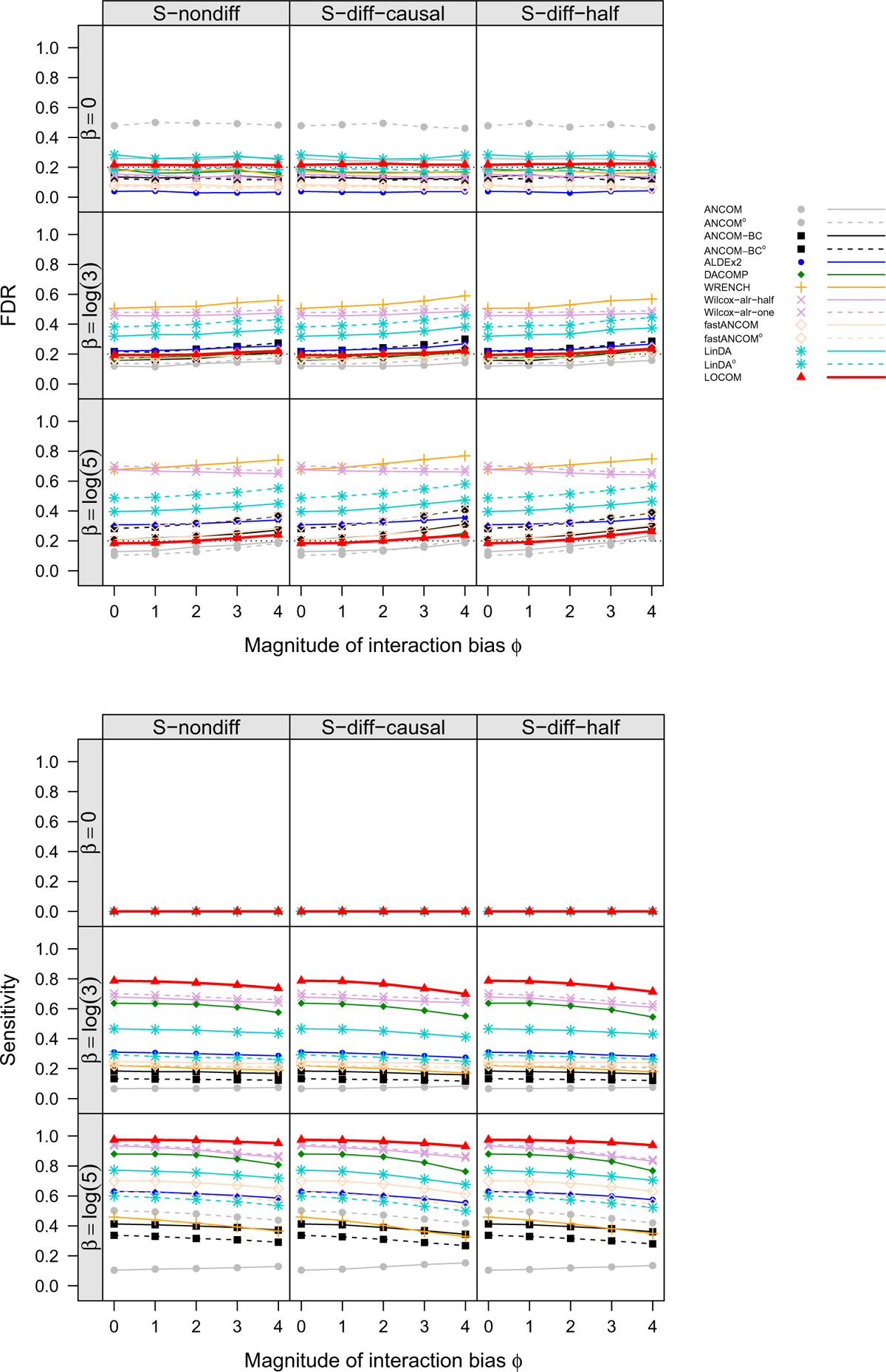
FDR and sensitivity at the nominal FDR 20% (black dotted line) of LOCOM and other compositional methods for data simulated under M1 (randomly selecting 20 taxa to be causal with effect size *β*) and had a binary trait. With superscript ^*o*^, the filter for rare taxa that are present in less than 10% of samples as adopted by the original programs was used; without ^*o*^, the more stringent filter with 20% cutoff as recommended by LOCOM was used. Details of Wilcox-alr-half and Wilcox-alr-one are found in [2]. All results were based on 1000 replicates of data.

**Figure 2:**
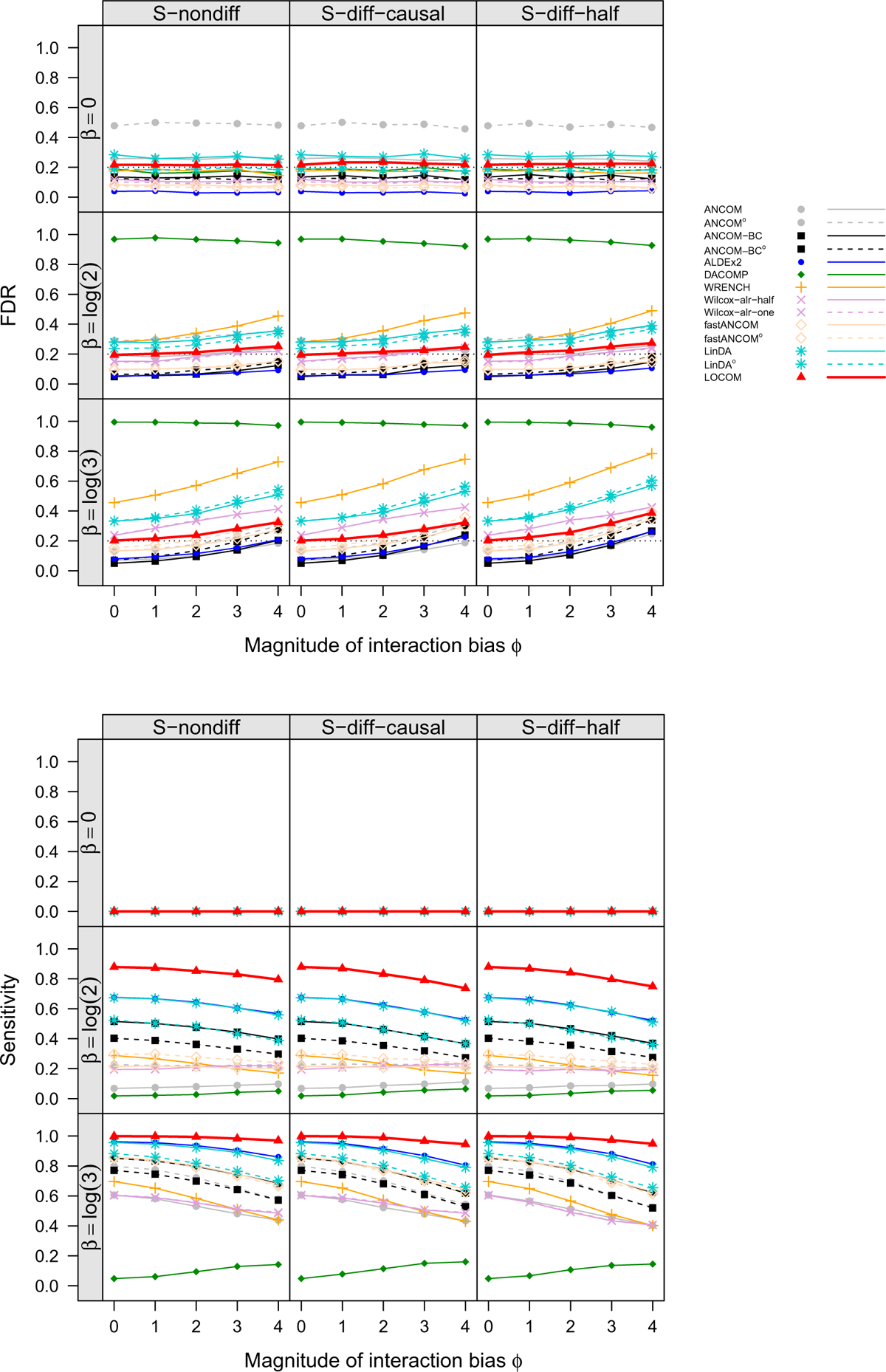
Same as the caption to Figure 1 except that the data were simulated under M2 (choosing the top five most abundant taxa to be causal with effect size *β*).

The other methods had very different performance even when there were only main effect biases (*ϕ* = 0). ANCOM-BC and fastANCOM performed the best among the other methods, controlling FDR for the full range of *β* values we have considered when *ϕ* was small and having similar FDR inflation as LOCOM when both *β* and *ϕ* became large; however, this performance came at the cost of a substantially lower sensitivity than LOCOM. ANCOM had moderately inflated FDR when *β* was small but performed better when *β* was increased; nevertheless, its sensitivity was among the lowest. ALDEx2 had inflated FDR and poor sensitivity when *β* was large, in M1 but not in M2. WRENCH, Wilcox-alr-half, and Wilcox-alr-one had highly inflated FDR except in the global null case and LinDA had highly inflated FDR in all cases including the global null. These results were based on data after applying the filter of rare taxa as recommended by LOCOM; in general, ANCOM, ANCOM-BC, fastANCOM and LinDA had worse FDR control when a less stringent filter was applied. Note that, unlike the LOCOM paper that used the older version (v1.1) of DACOMP, we used the latest version (v1.26) here, which yielded highly inflated FDR and low sensitivity in M2 (because some reference taxa were incorrectly selected to be causal taxa).

## Discussion

In this study, we found that LOCOM was robust to not only main effect biases but also a reasonable range of interaction biases. The other methods tended to have inflated FDR even when there were only main effect biases; many of them did not control the FDR even when there was no experimental bias at all (results shown in [2]). LOCOM maintained the highest sensitivity among all methods even when the other methods did not control the FDR. Therefore, we conclude that LOCOM outperforms most (if not all) existing methods for compositional analysis of microbiome data.

The robust performance of all methods to the interaction bias is likely because each term 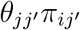 in (1) that governs the effect of taxon *j′* on the bias factor of taxon *j* is small because the relative abundance *π_ij′_* is generally small. Further, even if all contributions from different taxa to the total interaction bias 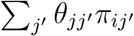 have the same sign, any non-zero mean interaction bias across taxa (or individuals) would be automatically included in the normalization factor *α_i_* (or the main effect *γ_j_*); Thus, the interaction bias is “automatically centered” constraints, which may also decrease the apparent effect of the interaction bias.

We used the Brooks mock community data to motivate our simulation studies, which may have two limitations. First, the Brooks samples contain at most seven taxa, all having equal true relative abundance; other mock community datasets, especially datasets that contain a large number of taxa and that mimic real microbiome data, should be used to study the interaction bias. Second, bacteria in a real microbial community may have different interactions from bacteria in a mock community. Therefore, study of experimental biases and their impact on downstream analysis continues to be an important topic in the foreseeable future.

## Acknowledgments

This research was supported by the National Institutes of Health awards R01GM116065 (Hu, Satten) and R01GM141074 (Hu, Satten).

