## Supplementary Materials for "Impact of experimental bias on compositional analysis of microbiome data"

### Supporting Information

#### Motivation on the value of $\phi$ and the sign of (2)

We used the main effect and interaction biases in the seven taxa of the Brooks mock community samples that were estimated by Zhao and Satten [3] and presented in their Table 4. Note that the interaction biases in [3] depend on the presence-absence status (0/1) of the other taxa, rather than their relative abundances as we modeled in (1), so the "interaction effect" in Table 4 of [3] corresponds to the product $\theta_{jj'}\pi_{ij'}$ in our model (1). Because the Brooks data are dominated by samples that have three taxa with equal proportions, i.e., $\pi_{ij'} = 1/3$, and the mean of the "interaction effect"-"main effect" ratios in Table 4 of [3] is 0.204, we obtained an estimate for our $\theta_{jj'}/\gamma_j$ to be 0.612. In our simulations, we varied the value of $\phi$ from 0 to 4, which includes our empirical estimate 0.612 while also extending the range of $\phi$ to allow stronger interaction biases than found in the Brooks data. Table 4 of [3] also shows that the interaction bias in taxon $j$ caused by taxon $j'$ has an opposite sign to the sign of the main effect bias in taxon $j'$, except for some very small interaction biases.

#### Details of the simulation settings

In M1, we randomly sampled 20 taxa to be causal from the set of taxa having mean relative abundances in the upper-respiratory-tract (URT) microbiome data [11] greater than 0.005 (but excluding the most prevalent taxon). In M2, we selected the top five most abundant taxa (having mean relative abundances 0.105, 0.062, 0.054, 0.050, and 0.049) to be causal. For simulations with a binary confounder, we assumed that the confounder was associated with 20 taxa under M1 (10 sampled at random from the causal taxa and 10 from the null taxa) and 5 taxa under M2 (2 from the causal taxa and 3 from the null taxa). We set the sample size to 100. Let $T_i$ denote the trait and $C_i$ the confounder for the $i$th sample so that $X_i = (T_i, C_i)^{\mathrm{T}}$. To generate a binary trait, we selected an equal number of samples with $T_i = 1$ and $T_i = 0$. When a binary confounder was present, we drew $C_i$ from $Bernoulli(0.2)$ in samples with $T_i = 0$ and from $Bernoulli(0.8)$ in samples with $T_i = 1$. To generate a continuous trait, we sampled $T_i$ from $U[-1, 1]$. To simulate read count data for the 856 taxa, we first sampled the baseline relative abundances $\pi_i^{(0)} = \left(\pi_{i1}^{(0)}, \pi_{i2}^{(0)}, \ldots, \pi_{iJ}^{(0)}\right)$ of all taxa for each sample from $Dirichlet(\overline{\pi}, \theta)$, in which $\overline{\pi}$ and $\theta$ took the estimated mean and overdispersion (0.02) from fitting the Dirichlet-Multinomial model to the URT data. We formed the *true* relative abundances $\pi_{ij}$ for all taxa by spiking causal taxon $j'$ with an $\exp(\beta_{j',1}T_i)$ fold change, spiking confounder-associated taxon $j''$ with an $\exp(\beta_{j'',2}C_i)$ fold change, and normalizing the relative abundances, so that

$$\pi_{ij} = \frac{\exp\left(\beta_{j,1}T_i + \beta_{j,2}C_i\right)\pi_{ij}^{(0)}}{\sum_{j^*=1}^{J}\exp\left(\beta_{j^*,1}T_i + \beta_{j^*,2}C_i\right)\pi_{ij^*}^{(0)}} \ .$$

We formed the *observed* relative abundances $p_{ij}$ by additionally multiplying the bias factor $\exp(\gamma_j + \sum_{j'}\theta_{jj'}\pi_{ij'})$, so that

$$p_{ij} = \frac{\exp\left(\gamma_j + \sum_{j'}\theta_{jj'}\pi_{ij'} + \beta_{j,1}T_i + \beta_{j,2}C_i\right)\pi_{ij}^{(0)}}{\sum_{j^*=1}^{J}\exp\left(\gamma_{j^*} + \sum_{j'}\theta_{j^*j'}\pi_{ij'} + \beta_{j^*,1}T_i + \beta_{j^*,2}C_i\right)\pi_{ij^*}^{(0)}} \ .$$

Note that $\beta_{j,1} = 0$ for null taxa and $\beta_{j,2} = 0$ for confounder-independent taxa. For simplicity, we set $\beta_{j,1} = \beta$ for all causal taxa. We generated the main effect bias $\gamma_j$ from $N(0, 0.8^2)$, which gave a range between 0.2 and 5 for most (95%) fold changes ($\exp\gamma_j$) caused by the main effect bias. The scheme for generating the interaction bias $\theta_{jj'}$ are described below. Finally, we generated the taxon count data using the Multinomial model with mean $(p_{i1}, p_{i2}, \ldots, p_{iJ})$ and library size sampled from $N(10000, (10000/3)^2)$ and left-truncated at 2000.

To generate the interaction bias $\theta_{jj'}$, it remained to generate the error term $\epsilon_{jj'}$. In S-nondiff, we sampled $\epsilon_{jj'}/2$ from $N(0.5, 0.1^2)$ for all taxa pairs $j$ and $j'$. In S-diff-causal, we modified S-nondiff to sample $\epsilon_{jj'}/2$ from $Beta(0.5, 0.5)$ when taxon $j$ was causal (and for all $j'$). Both distributions have mean 0.5; however, $N(0.5, 0.1^2)$ has one mode at 0.5 and $Beta(0.5, 0.5)$ has two modes 0 and 1. In S-diff-half, we used $N(0.5, 0.1^2)$ for half of randomly selected taxa $j$s and $Beta(0.5, 0.5)$ for the remaining half of taxa. The distribution (figures not shown) of the $j$th bias factor due to taxon-taxon interactions, $\eta_{ij} = \sum_{j'}\theta_{jj'}\pi_{ij'}$, across samples has the following features. In all scenarios and for all taxa $j$s, the mean of $\eta_{ij}$ was approximately zero due to the averaging of contributions with different directionalities. The variance of $\eta_{ij}$ increased as $\gamma_j$ increased at a given $\phi$. S-nondiff and S-diff-causal had the same

$\eta_{ij}$ values at the null taxa and different $\eta_{ij}$ values at the causal taxa. Finally, in S-diff-half, $\eta_{ij}$ had larger variance in one half of taxa than the other half of taxa.
